# A Combined approach of MALDI-TOF Mass Spectrometry and multivariate analysis as a potential tool for the detection of SARS-CoV-2 virus in nasopharyngeal swabs

**DOI:** 10.1101/2020.05.07.082925

**Authors:** María Florencia Rocca, Jonathan Cristian Zintgraff, María Elena Dattero, Leonardo Silva Santos, Martín Ledesma, Carlos Vay, Mónica Prieto, Estefanía Benedetti, Martín Avaro, Mara Russo, Fabiane Manke Nachtigall, Elsa Baumeister

**Author notes:** These authors contributed equally to this work (María Florencia Rocca, Jonathan Cristian Zintgraff, María Elena Dattero). **Corresponding author:** María Florencia Rocca.

## Abstract

Coronavirus disease 2019 (COVID-19) is caused by the severe acute respiratory syndrome coronavirus 2 (SARS-CoV-2). The rapid, sensitive and specific diagnosis of SARS-CoV-2 by fast and unambiguous testing is widely recognized to be critical in responding to the ongoing outbreak. Since the current testing capacity of RT-PCR-based methods is being challenged due to the extraordinary demand of supplies, such as RNA extraction kits and PCR reagents worldwide, alternative and/or complementary testing assays should be developed. Here, we exploit the potential of mass spectrometry technology combined with machine learning algorithms as an alternative fast tool for SARS-CoV-2 detection from nasopharyngeal swabs samples. According to our preliminary results, mass spectrometry-based methods combined with multivariate analysis showed an interesting potential as a complementary diagnostic tool and further steps should be focused on sample preparation protocols and the improvement of the technology applied.

## Introduction

The novel coronavirus disease 2019 (COVID-19), caused by the SARS-CoV-2 virus, was declared a pandemic by the World Health Organization on March 12^th^ 2020 following its emergence in Wuhan China. As of the April 4^th^ 2020 there were over 1.2M confirmed cases of COVID-19 in 175 countries, with over 65,000 fatalities (Johns Hopkins Coronavirus Resource Center, 2020). SARS-CoV-2 is one of four new pathogenic viruses which have jumped from animal to human hosts over the past 20 years, and the current pandemic sends warning signs about the need for preparedness and associated research. Clinical presentation of COVID-19 ranges from mild to severe with a high proportion of the population having no symptoms yet being equally infectious (Yang et al., 2020; Zhou et al., 2019). Together, these features have led to intensive lockdown measures in most countries with the aim to restrict the spread of the virus, limit the burden on healthcare systems and reduce mortality rate. In parallel, there has been an extraordinary response from the scientific community. These collective efforts aim to understand the pathogenesis of the disease, to evaluate treatment strategies and to develop a vaccine at unprecedented speeds in order to minimize its impact on individuals and on the global economy (Li, 2016; Wenzhong and Hualan, Preprint).

The rapid, sensitive and specific diagnosis of SARS-CoV-2 by fast and unambiguous testing is widely recognized to be critical in responding to the outbreak. Since the current testing capacity of RT-PCR-based methods is being challenged due to the extraordinary global demand of supplies such as RNA extraction kits and PCR reagents, alternative and/or complementary testing assays need to be deployed now in an effort to accelerate our understanding of COVID-19 disease (Chin et al., 2020; Antezack et al., 2020).

The aim of this work was to assess the potential of MALDI-TOF MS technology to create mass spectra from nasopharyngeal swabs in order to find specific discriminatory peaks by using machine learning algorithms, and whether those peaks were able to differentiate COVID-19 positive samples from COVID-19 negative samples.

## Materials and methods

### Sample preparation and MALDI-TOF data acquisition

#### Samples

First, we analyzed in triplicate 25 samples of nasopharyngeal swab, preliminary tested by RT-PCR (Corman et al., 2020) at the Reference Respiratory Virus Laboratory INEI-ANLIS “Dr. Carlos G. Malbrán” in Argentina. Of the 25 samples, 13 were positive for SARS-CoV-2 and 12 had nondetectable viral load for SARS-CoV-2 (8/12 were positive for the other respiratory virus such as respiratory syncytial virus, Measles virus, Influenza A and B virus, and endemic human coronavirus).

The analysis was carried out using the Bruker Daltonics MicroFlex LT instrument version 3.4 (Bruker Daltonics, Bremen, Germany). Briefly, for the assay, 1ul of each nasopharyngeal swab was applied onto 3 wells of the steel MALDI plate (MSP 96 target ground steel; Bruker Daltonics), after the wells have got dried, they were overlaid with 1 μl of HCCA matrix (a solution containing α-cyano-4-hydroxycinnamic acid diluted into 500μL of acetonitrile, 250μL of 10% trifluoroacetic acid and 250μL of HPLC grade water). All manipulations were performed under certified class II biological safety cabinet TELSTARTM BIO IIA (Thermo Fischer Scientific, Villebon sur Yvette, France)and wearing all the appropriate personal protective equipment (PPE) required to comply with biosafety standards (World Health Organization, 2020). After drying for a few minutes at room temperature, the plate was loaded into the instrument and analyzed using MALDI-TOF software.

#### Spectra acquisition

Spectra were acquired manually (mode OFF), reaching 160-200 laser shots per well and within a mass-range of 1960-20200 Da using FlexControl software v3.4 (Bruker Daltonics, Bremen, Germany). The platform was previously calibrated using the Bruker Daltonics Bacterial Test Standard.

All spectra were verified using the Flex Analysis v3.4 software (Bruker Daltonics, Bremen, Germany). The spectra selected for model generation and classification were treated according to a standard workflow including the following steps: baseline subtraction, normalization, recalibration, average spectra calculation, average peak list calculation, peak calculation in the individual spectra and normalization of peak list for model generation.

## MALDI-TOF MS spectra analysis

### Database development. MSP library construction

Main Spectrum Profiles (MSPs) were performed according to manufacturer’s instructions. All 25 samples were used to build an “in-house” database with Maldi Biotyper OC V3.1 (Bruker Daltonics, Bremen, Germany); in addition, dendrograms were performed to assess the relatedness of these MSPs using default settings.

### Biomarker assignment

Spectra files from MSPs were exported as mzXML files using CompassXport CXP3.0.5. (Bruker Daltonics, Bremen, Germany) for visual analysis. At the same time, a new database was created in Bionumerics v7.6.2 (Applied Maths, Ghent, Belgium) according to the manufacturers’ instructions. All raw spectra were imported into the Bionumerics database with x-axis trimming to a minimum of 1960 m/z.

Baseline subtraction (with a rolling disc with a size of 50 points), noise computing (continuous wavelet transformation, CWT), smooth (Kaiser Window with a window size of 20 points and beta of 10 points), and peak detection (CWT with a minimum signal to noise ratio of 1) were performed. Spectrum summarizing, peak matching and peak assignment was performed according to instructions from Bionumerics. In short, all raw spectra were summarized into isolate spectra, and peak matching, using the option “existing peak classes only”, was performed on isolate spectra using a constant tolerance of 1.9, a linear tolerance of 550 and a peak detection rate of 10%. Binary peak matching tables were exported to summarize the presence of peak classes. By assigning biomarkers, only the presence and absence of peaks was investigated. To also assess quantitative peak data such as peak intensity and peak area, a multivariate unsupervised statistical tool PCA was additionally performed. All samples were examined for the unique masses (± 10 Da).

### Classifier models based on machine learning

#### ClinProTools software

Peak data of all samples were used to define and train machine learning based classifiers using ClinProTools V.2.2 software (Bruker Daltonics, Bremen, Germany) according to the manufacturer’s instruction. In brief, data analysis began with the loading of the raw data into the software and they were grouped into different classes; first we created a two-class model (Class 1 = SARS-CoV-2 detectable samples; Class 2= undetectable viral load samples). The distribution of the elements is showed in **Fig. 1**.

**Figure 1.**
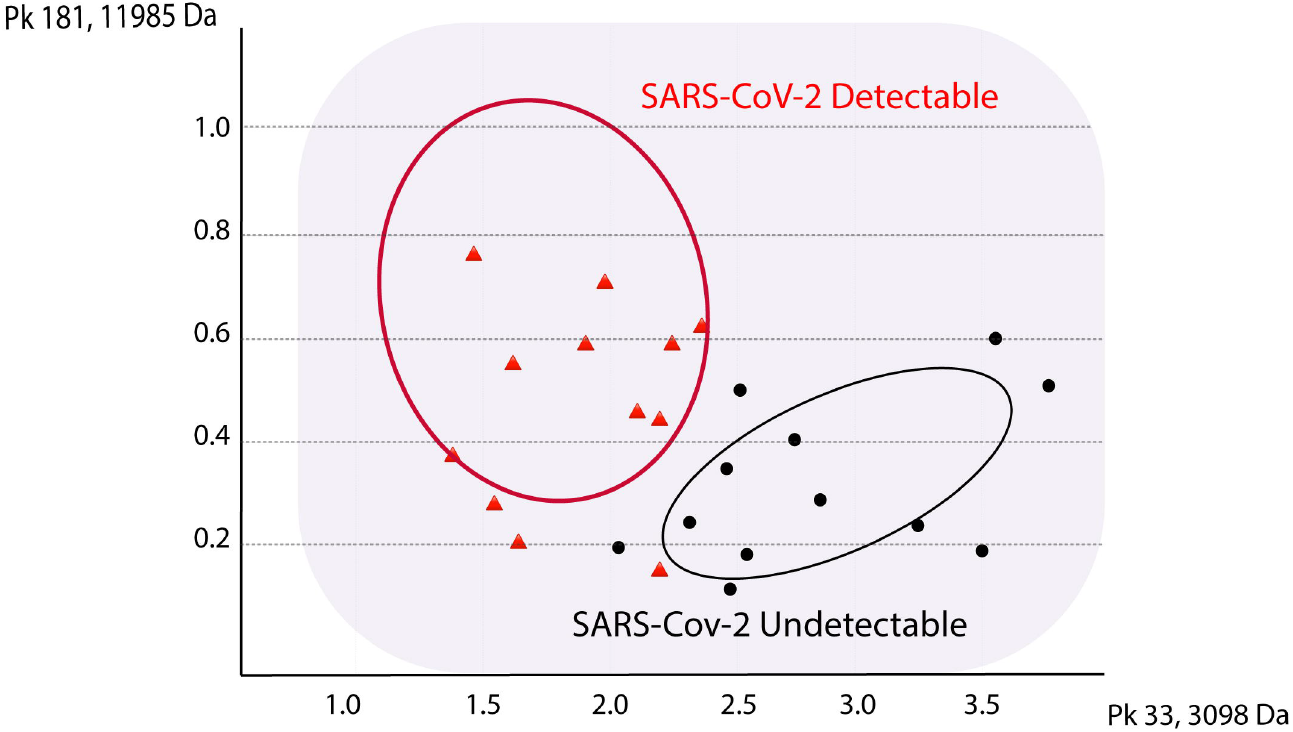
2D Peak Distribution Plot of 2 classes model. This plot displays the distribution of two selected peaks in the non-excluded spectra on the loaded model generation classes. The data is shown on a two-dimensional plane. By default, the first two (=best separating) peaks of the current statistic sort order are displayed. The ellipses represent the standard deviation of the peak area/intensities.

Then a three-class model (Class 1 = SARS-CoV-2 detectable samples; Class 2 = undetectable viral load samples and Class 3 = other respiratory virus) was developed. Pretreatment, normalization, baseline subtraction, peak defining (range 1960-20000 m/z), recalibration; and then, the automatic comparison of multiple spectra was performed. Values of m/z from the average spectra of each class and informative peaks were identified according to their statistical significance, as determined by the different statistical tests supported by ClinProTools: Anderson-Darling test, t-/ analysis of variance (ANOVA) test and Wilcoxon/Kruskal-Wallis tests. Informative peaks were those showing a significant difference between the classes whether: the p-value for the Anderson–Darling test was >0.05 and for the t-/ANOVA or Wilcoxon/Kruskal–Wallis test was ≤0.05, or if the p-value for the Anderson–Darling test was ≤0.05 and for the Wilcoxon/ Kruskal–Wallis test was ≤0.05 (Stephens, 1974). The best top ten peaks are summarized in **Table 1**.

**Table 1.** Characteristic MALDI-TOF MS peaks, obtained by ClinProTools software.

| 2 Classes model peak statistic table. |  |  |  |  |  |
| --- | --- | --- | --- | --- | --- |
| Index | Mass | DAve | PTTA | PWKW | PAD |
| 33 | 3098.4 | 1.21 | 0.00084 | 0.00025 | 0.198 |
| 181 | 11985.13 | 0.61 | 0.00169 | 0.00183 | < 0.000001 |
| 34 | 3142.88 | 2.08 | 0.00226 | 0.00000558 | 0.000742 |
| 19 | 2435.27 | 1.46 | 0.00226 | 0.00487 | 0.372 |
| 57 | 4035.25 | 1.45 | 0.00226 | 0.0014 | 0.0000118 |
| 124 | 6825.04 | 0.74 | 0.00226 | 0.000216 | < 0.000001 |
| 175 | 11370.78 | 0.29 | 0.00265 | 0.00646 | < 0.000001 |
| 56 | 3937.83 | 1.2 | 0.00411 | 0.000216 | 0.000815 |
| 114 | 5993.37 | 0.64 | 0.00966 | 0.0184 | < 0.000001 |
| 26 | 2633.7 | 2.83 | 0.0106 | 0.000216 | 0.00000165 |

| 3 Classes model peak statistic table. |  |  |  |  |  |
| --- | --- | --- | --- | --- | --- |
| Index | Mass | DAve | PTTA | PWKW | PAD |
| 9 | 2178.68 | 7.24 | < 0.000001 | 0.0000104 | < 0.000001 |
| 10 | 2200.65 | 2.95 | < 0.000001 | 0.000154 | < 0.000001 |
| 11 | 2223.56 | 2.72 | < 0.000001 | 0.0000378 | < 0.000001 |
| 62 | 4470.31 | 1.51 | < 0.000001 | 0.0000135 | < 0.000001 |
| 14 | 2340.85 | 4.29 | 0.000011 | 0.0000104 | 0.0000314 |
| 29 | 3319.23 | 3.23 | 0.0000268 | 0.0000104 | < 0.000001 |
| 105 | 6177.9 | 0.8 | 0.0000268 | 0.0000104 | < 0.000001 |
| 64 | 4573.76 | 3.71 | 0.000038 | 0.0000104 | < 0.000001 |
| 155 | 10095.58 | 0.41 | 0.000038 | 0.0000183 | 0.0000492 |
| 5 | 2118.03 | 3.93 | 0.0000481 | 0.000121 | 0.00428 |

DAve: Difference between the maximal and the minimal average peak intensity of all classes.
PTTA: P-value obtained through t-test. PWKW: P-value obtained through Wilcoxon/Kruskal-Wallis test. PAD: P-value obtained through Anderson-Darling test.

Classification models were generated using all three available algorithms (Supervised Neural Network, Genetic Algorithm and QuickClassifier) and they were later compared. For each model, the recognition capability (RC) and cross validation (CV) percentage was generated to demonstrate the reliability and accuracy of the model. RC and CV percentages were indicators of the model’s performance and useful predictors of the model’s ability to classify test samples. The model with the highest RC and CV values was used in the analysis.

### Statistical analysis

For evaluating the performance of the different approaches mentioned, accuracy, sensitivity, specificity, positive prediction and negative prediction were calculated (ClinPro Tools 3.0: User Manual, 2011).

## Results

The results were analyzed according to the following approaches: Biomarker findings, construction of an “in-house” database and through the design of predictive models by machine learning.

### Detection of potential Biomarkers

Manual analysis of the spectra obtained by Flex Analysis v3.4 software revealed one potential peak of negativity which was not detected in most of the positive samples: **4551 Da.**

When BioNumerics software was used, another potential biomarker was found in 60% of negative samples and only in 16% of positives: **3142 Da.**

This peak is also detected with reproducible intensity in the average spectrum of that same class in ClinProTools **(Fig. 2).**

**Figure 2.**
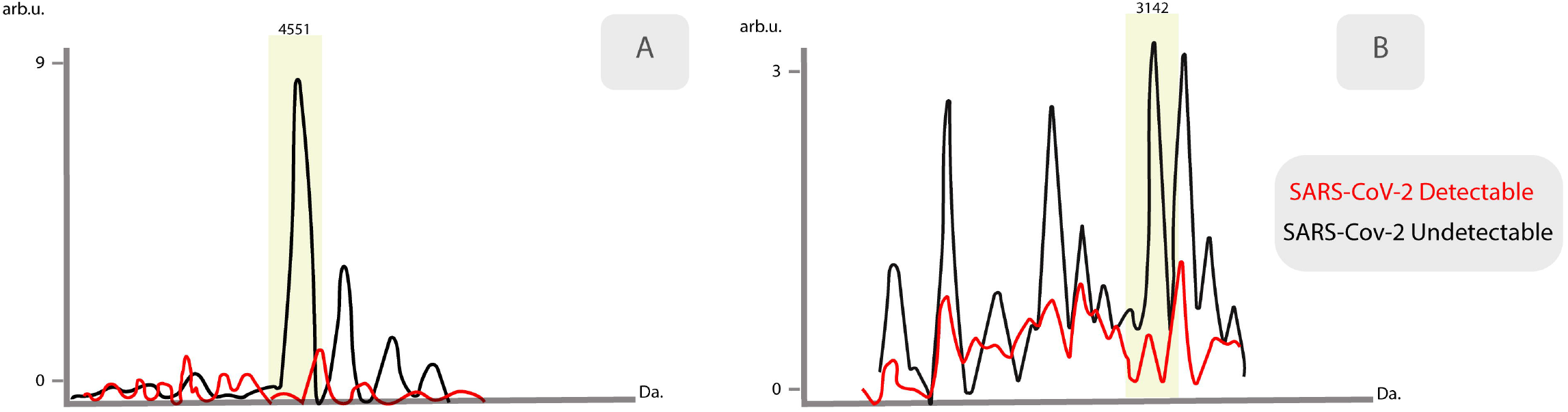
Average spectra of characteristic peaks among detectable viral load samples versus undetectable viral load samples. Intensities of characteristic peaks: A- (*m/z* 4551) Obtained by Flex Analysis v3.4 software manual analysis. B- (*m/z* 3142) Obtained by Bionumerics v7.6.2 software.

### Evaluation of the novel “in-house” database

The novel database was challenged with 30 previously-characterized samples different than those used to create it. They were processed in the same way as to create the complementary library and the wells were read in the MALDI Biotyper OC v3.1 software according to the manufacturer’s recommendations, for an offline classification, obtaining the following results: 12 of 19 positive samples (63%) were correctly identified with score values >1.70; 1 of 19 (5%) was miss identified and 6 of 19 (32%) presented low score values, making identification not reliable. Contrasting, 8 of 11 of negative samples (73%) were identified correctly presenting score values above 1.70; 2 samples (18%) were miss identified and only 1 (9%) presented low score values. Samples with low score values were not included in the statistical analysis.

### Machine Learning models

The results of Recognition capacity (RC) and Cross-Validation (CV) values of all models used are summarized in Table 2.

**Table 2.** Machine Learning results.

| Classifier Models | Recognition Capacity (%) | Cross Validation (%) |
| --- | --- | --- |
| Genetic Algorithm<br>2 classes | 100 | 93.9 |
| Supervised Neural<br>Network<br>2 classes | 100 | 82.9 |
| QuickClassifier<br>2 classes | 82.1 | 68.1 |
| Genetic Algorithm<br>3 classes | 100 | 85.5 |

With the application of a combined algorithm (GA 3 classes/ GA 2 classes/ SNN 2 classes, in that order) and considering two of three models as concordant, it was possible to identify 68% (13/19) SARS-CoV-2 positive samples, on the other hand, 16% (3/19) were miss identified and 3 of 19 (16%) were discordant for the GA 2 classes and SNN 2 classes but correctly identified by the GA 3 classes (not included in the statistical analysis).

Considering the SARS-CoV-2 undetectable samples: 91% (10/11) were correctly identified and only 1 sample was miss classified **(Table 3).**

**Table 3.** Parameters of the different approaches evaluated.

| Methods Evaluated | Accuracy (%) | Specificity (%) | Sensitivity (%) | Positive Prediction (%) | Negative Prediction (%) |
| --- | --- | --- | --- | --- | --- |
| "in-house" Database | 86.9 | 80.0 | 92.3 | 85.7 | 88.9 |
| Machine Learning | 85.2 | 90.9 | 81.2 | 92.8 | 76.9 |

## Discussion

MALDI-TOF MS is a simple, inexpensive and fast technique that analyses protein profiles with a high reliability rate, and could be used as a rapid screening method in a large population (Croxatto et al., 2012). These preliminary results suggest that MALDI-TOF MS coupled with ClinProTools software represents an interesting alternative as a screening tool for diagnosis of SARS-CoV-2, especially because of the good performance and accuracy obtained with samples in which viral presence was not detected. More samples need to be analyzed in order to make a definitive statement. However, this study using MALDI-TOF combined with machine learning has proven to be, as far as we know, a revolutionary alternative that deserves further development.

## Conclusions

The identification of specific biomarkers responsible for each peak or group of peaks represents a difficult and demanding task that requires further specific studies. Based on the promising preliminary results, we should focus on the improvement of this potential diagnosis approach by assaying various techniques for proteins extraction to the clinical samples and on the expansion of the complementary database in the near future.

These results constitute the basis for further research and we encourage researchers to explore the potential of MALDI-TOF MS in order to assess the feasibility of this technology, widely available in clinical microbiology laboratories, as a fast and inexpensive SARS-CoV-2 diagnostic tool.

